# High intraspecific growth variability despite strong evolutionary heritage in a neotropical forest

**DOI:** 10.1101/2022.07.27.501745

**Authors:** Sylvain Schmitt, Bruno Hérault, Géraldine Derroire

**Author notes:** Correspondence: Sylvain Schmitt, UMR EcoFoG, Campus Agronomique, 97310 Kourou, French Guiana.

## Abstract

Individual tree growth is a key determinant of species performance and a driver of forest dynamics and composition. Previous studies on tree growth unravelled the variation in species growth as a function of demographic trade-offs that are partially predicted using functional traits. They have explored the environmental determinants of species growth potential and the variation of intraspecific growth over space and time due to environment and biotic factors. However, variation in individual growth within species remains underexplored for a whole community and the relative role of species’ evolutionary heritage and of local environments remains unquantified. Here, based on 36 years of diameter records for thousands of mapped individuals belonging to 138 species, we assessed individual tree growth potential in a local neotropical forest community in the Amazon basin. We further related variation in individual growth potential with taxonomic levels, local topography, and neighbourhood crowding, before exploring species growth potential link to functional traits and distribution along the phylogeny. We found that most of the variation in growth potential was individual, and that taxonomic structure explained a third of the observed variation. Species growth potential was phylogenetically conserved with positive conservatism up to the genus level in the vast majority of species. Functional traits of roots, wood and leaves together predicted species growth potential. Phylogeny suggested joint selection of species’ growth strategies and associated functional traits during convergent evolutions. Finally, neighbourhood crowding had a significant effect on individual growth potential, although much of this inter-individual variation remains largely unexplained and the underlying ecological and evolutionary factors are still little explored. The high intraspecific variation observed could allow individuals in these hyperdiverse ecosystems to respond to the variable light and competitive conditions offered by successional niches during forest gap dynamics.

## Introduction

Individual growth is a key determinant of species performance (Violle et al., 2007) and a driver of forest dynamics (Hérault et al., 2010) and composition (Russo et al., 2008). Thus, a comprehensive understanding of the determinants of tree growth is of primary importance for predicting the fate of tropical forests, especially in the face of increasing anthropogenic disturbances. Tree growth plays a major role in the ecological strategies of tree species through demographic trade-offs (Rüger et al., 2019) including a growth-mortality trade-off (Aubry-Kientz et al., 2015; Philipson et al., 2014; Wright et al., 2010). This trade-off opposes fast-growing species to slow-growing species (Aubry-Kientz et al., 2015; Phillipson et al., 2014;), with lower support costs but at greater risk of damage and increased mortality (King et al., 2006). To forecast the dynamics of tropical forests, efforts have been made to predict the growth of species. To this end, functional traits have been widely explored and used (Hérault et al., 2011, Visser et al., 2016, Osazuwa-Peters et al., 2017, Poorter et al., 2010). Functional traits are defined as phenotypic traits that have an impact on fitness through their effect on individual performance, which is defined as the ability to recruit, grow, survive and reproduce (Violle et al. 2007). Functional traits are therefore expected to play a role in species growth (but see Yang et al., 2018). For instance, wood density has been shown to be an important determinant of species growth potential (King et al., 2005; King et al., 2006; Hérault et al., 2011; Visser et al., 2016), as well as species maximum diameter and height, δ^13^*C* of leaf (Hérault et al., 2011), wood anatomical features (Osazuwa-Peters et al., 2017; Poorter et al., 2010) and hydraulic conductance (Poorter et al., 2010).

Multiple environmental determinants of species growth have already been identified, which in turn affect species distribution and shape community structure. Light interception was recognised early on as a key factor in the growth of species (King et al., 2005). The role of light has notably been demonstrated experimentally in seedlings with species showing a trade-off between shade tolerance and establishment in gaps (Baraloto et al., 2005). Fast-growing species are found in environments with high access to light, where shade-tolerant species show reduced growth (Wright et al., 2010). Treefalls open gaps in the forest creating bright environments and reduced competition from large trees. During forest succession, gaps are filled by vegetation, resulting in shadier environments and increased competition from large trees. This process, called forest gap dynamics (Martinez-Ramos *et al.,* 1989), produces successional niches, which favour species with a variety of survival and growth strategies (Hérault *et al.,* 2010; Rüger *et al.,* 2009). Fast-growing pioneer species quickly colonise treefall gaps, while slower-growing late-successional species gradually establish themselves in shadier environments (Craven *et al.,* 2015). Species distribution is also driven by soil (Kupers et al., 2019). Nutrient availability together with water shape species performance with fast-growing species dying more on the poorest habitat and slow-growing species being outcompeted in resource-rich habitats (Russo et al., 2008).

Species strategies of resource use, and the functional traits used to study these strategies, are however still poor predictors of individual tree rates (Yang *et al*., 2018), partly because they ignore individual variation, and because critical aspects of demographic rates are not captured by most measured functional traits. Individual- and species-based approaches to functional traits are conceptually fundamentally different (Poorter *et al*., 2018): the species-based approach focuses on potential traits and rates and the individual-based approach focuses on realised traits and rates. Individual growth rates depend on access to resources, which are modulated by biotic interactions with neighbouring trees. Two environmental drivers can capture these interacting effects. (1) Topography is driving both water and nutrient availability in tropical forests through the dissolution of iron oxides, litter- and tree-fall transfers and waterlogging (Ferry *et al*., 2010; John *et al*., 2007). Therefore, individual growth within species is faster in valleys than on ridges in neotropical forests (Fortunel *et al*., 2018; but see O’Brien and Escudero 2021). (2) Neighbourhood crowding through competition for resources, including light, nutrients and water, but also potentially through facilitation, is known to modulate tree growth (Uriarte et *al.,* 2004; Lewis and Tanner 2000). Individuals’ responses to neighbourhood crowding depend on the species identity of the focal tree and the composition of its neighbours (Uriarte et al., 2004; Lewis and Tanner 2000). Neighbourhoods interact with other abiotic factors, e.g. growth is faster in bottomland than on higher ground, but neighbourhood crowding has a stronger negative effect on growth in bottomland than in higher ground for one in ten species (Fortunel *et al.,* 2018). Neighbourhood crowding also captures the effect of forest gap dynamics, which is a key factor shaping the growth of individuals within species with increased tree growth near canopy openings (Hérault *et al*., 2010; Schmitt *et al.,* 2022).

Most of the studies of intraspecific variation in growth were carried out for a limited number of species of a community, and the consistency of the observed patterns across species of a community remains underexplored. This prevents assessing the relative contribution to individual growth variability of species evolutionary heritage (which we define here and throughout the manuscript as the contribution of phylogenetic relatedness to species phenotypic variation, following Souza *et al.,* 2016) linked to phylogenetic background and individual adaptation and plasticity in response to local environments. Individual trees however differ in many dimensions within and among species with genetic heritage and life history that may influence their growth response to local abiotic environments and biotic interactions (Le Bec et al., 2015). Genetic processes within and among species hold promises for understanding tree growth (Grattapaglia et al., 2009). Evolutionary heritage resulted in convergent species growth within tree genera from the Amazon Basin (de Souza *et al*., 2016). A recent study further suggested individual adaptation of tree growth within species to the successional niches generated by tropical forest gap dynamics in the tropical species from the *Symphonia* genus (Schmitt et al., 2022). The topographic origin of parent trees was also shown to determine the individual’s growth response of the offspring to increasing abundance of neighbours (O’Brien Escudero 2022). Altogether, these results plead for a better integration of species and individuals in tree growth studies, in order to better link individual tree growth to genetic, phylogenetic and environmental variations.

Here, we assessed tree growth potential for most individuals in a local neotropical forest community in the Amazon basin, and related variation in individual growth potential to taxonomic levels, local topography and neighbourhood crowding, in addition to the species phylogeny. Based on 36 years of diameter records for 7,961 of mapped individuals belonging to 138 species, we constructed individual ontogenetic growth trajectories to infer individual growth potential using a hierarchical Bayesian model. We used individual growth potential in a linear mixed model with topography, neighbourhood crowding and taxonomic level to explore the environmental drivers and evolutionary heritage of tree growth. We further explored the phylogenetic signal of species growth potential and the role of previously published functional traits in explaining species growth potential. We specifically addressed the following questions:

1. What is the importance of evolutionary heritage in the growth potential of tropical trees?
2. Can functional traits help predict species growth potential?
3. How important is the variability of individual growth potential and how does the local environment influence it?

We hypothesised species growth to show a phylogenetic signal with conserved interspecific growth in part influenced by species functional traits. We expected high intraspecific variation in tree growth influenced by topography and neighbourhood crowding.

## Material and Methods

### Study site

The study was conducted in the Guiana Shield, in the coastal region of French Guiana, at the Paracou field station (5°18′N, 52°53′W). The site is characterised by an average annual rainfall of 3041 mm and an average air temperature of 25.71°C (Aguilos et al. 2018). A rich tropical forest occupies this lowland area characterised by heterogeneous microtopographic conditions with numerous small hills generally not exceeding 45 m in altitude (Gourlet-Fleury et al. 2004). The site includes fifteen 6.25 ha plots and a 25 ha plot with trees mapped to the nearest metre and censused (diameter at breast height >10 cm) every 1 to 5 years since 1985. Nine of the plots were intentionally manipulated in 1986 with a range of disturbance intensities that created a variety of biotic environments (details of the experiment in Hérault and Piponiot 2018).

### Species and individuals

We focused on trees located 20 metres from any plot edge for neighbourhood analyses. We used only (i) trees recruited since the beginning of censuses to exclude large diameter trees that show little to no variation in their growth trajectories, (ii) trees with at least 10 measurements to better assess their complete growth trajectories and (iii) species with at least 10 trees meeting the previous requirements for a good representation of intraspecific variability. We did include disturbed plots in our study in order to explore a greater variety of neighbourhood crowding.

### Individual growth

We first used a reduced dataset to explore the best model shape (Tab. S1) and the best hierarchical integration of individual and species effects (Tab. S2) to infer individual growth potential. Based on goodness of fit (likelihood), cross-validation (leave-one-out estimate of the expected log pointwise predictive density, see Vehtari *et al*., 2007) and prediction quality (root mean square error of prediction), we chose to model tree diameter over time using the sum of the arithmetic series of the annual tree growth rate observed at the census year for each individual with a Gaussian distribution (model 1). At any year *t* since tree recruitment, individual annual growth rate *AGR_i,t_* can be defined following a Gompertz model (Hérault *et al*., 2011) based on individual diameter at breast height from previous census *DBH*_*i,t*−1_:

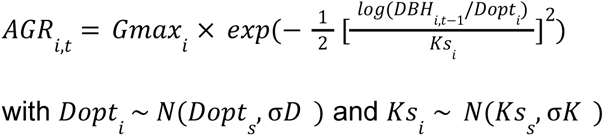

where *Gmax_i_* is the individual maximum growth potential, *Dopt_i_* is the optimal diameter at which the individual reaches its maximum growth potential, and *Ks_i_* is the kurtosis defining the width of the bell-shaped growth-trajectory (see figure 1 in Hérault *et al*., 2011). *Dopt_i_* and *Ks_i_* are random effects centred on species parameters *Dopt_s_* and *Ks_s_* with associated variances σ*D* and σ*K*. Thus, annual individual diameter *DBH_i,t_* can be calculated as previous year individual diameter *DBH*_*i,t*−1_ plus individual annual growth rate *AGR_i,t_*:

**Figure 1:**
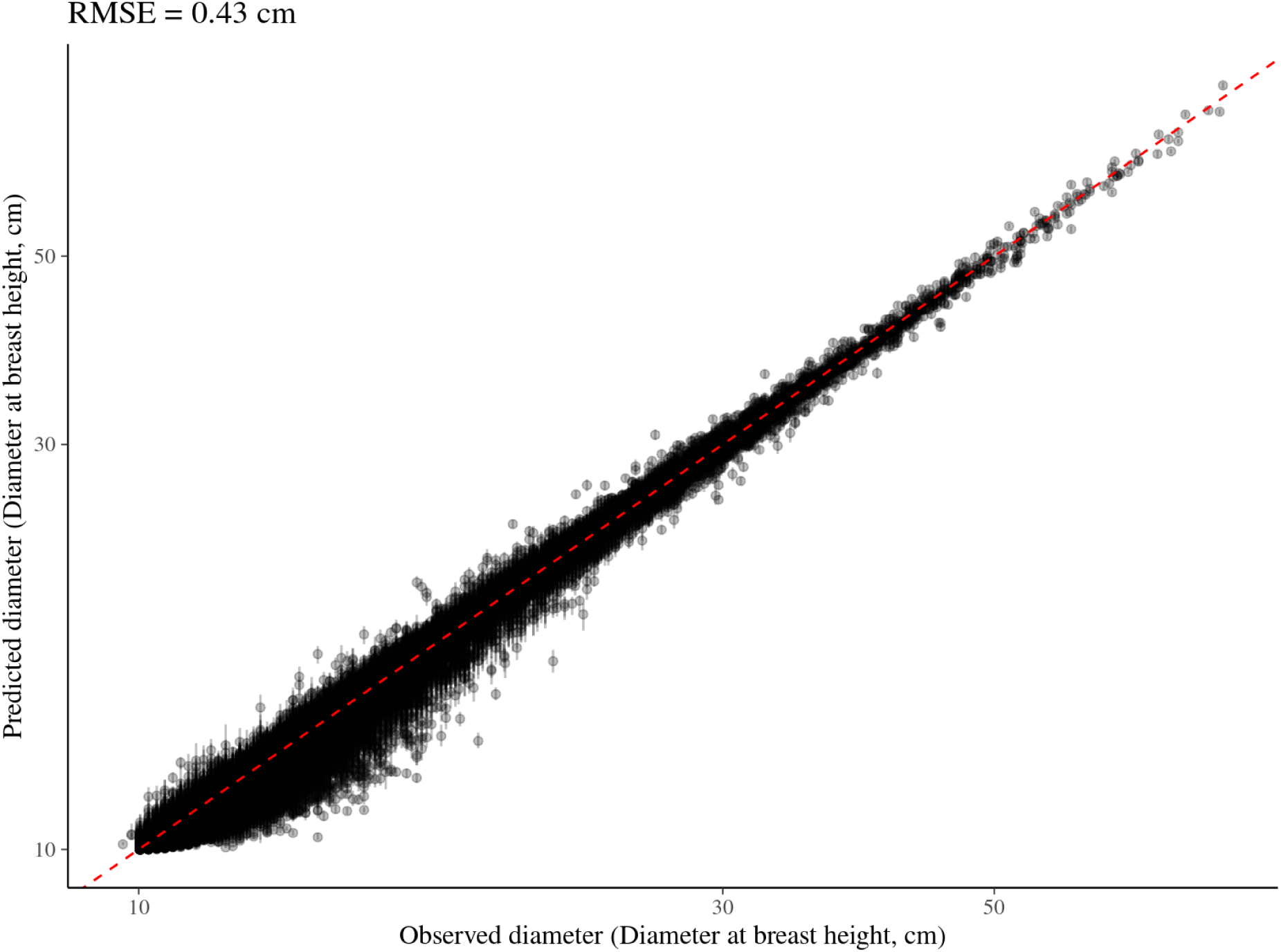
Goodness of fit for Bayesian inference of the individual growth model, illustrated by predicted versus observed diameters at breast height. The dots represent the median of the posterior distributions of predicted diameters while the error bars show the 90% credibility intervals. The red dashed line represents the expected value for a perfect prediction (1:1). The X and Y axes have been log-transformed to better represent the many small diameter values. An example of corresponding diameter trajectories can be found in Figure S4.

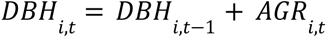

This arithmetic serie can be integrated from individual recruitment to build individual growth trajectory with annual individual diameter *DBH_i,t_*:

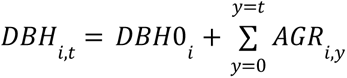

where *DBH0_i_* is individual diameter at recruitment. Finally, a model can be fitted to predict annual individual diameter *DBH_i,t_* with observed diameter from censuses using a Gaussian distribution:

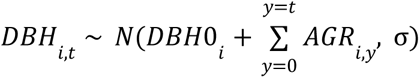

This model was fitted for every census *t* at which the individual was measured. A Bayesian method was employed to infer parameters using the stan language (Carpenter *et al*., 2017) and the *rstan* package (Stan Development Team, 2018) in the R environment (R Core Team, 2020).

### Descriptors of individual growth potential

We used the mean neighbourhood crowding index (NCI, Uriarte *et al.,* 2004) over the last 30 years, an indirect measurement of access to light and forest gap dynamics for each individual. The mean neighbourhood crowding index *NCI_i_* from tree individual i was calculated as follows:

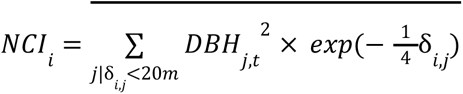

With *DBH_j,t_* the diameter of the neighbouring tree j in year t and δ_*i,j*_ its distance to the individual tree i. *NCI_i_* is computed for all neighbours at a distance δ_*i,j*_ inferior to the maximum neighbouring distance of 20 metres. The power of neighbours *DBH_j,t_* effect was set to 2 to represent an area. The decrease of neighbours’ diameter effect with distance was set to -¼ to represent trees at 20 metres of the focal trees having less than 1% of the effect of the same tree at 0 metres (*i.e.* exp(-¼ x 20) < 1%). *NCI_i_* is computed as the mean of *NCI_i,t_* per census over the last 30 years denoted by the overline in the equation.

We used the topographic wetness index (TWI) as proxy of the distribution of soil water and nutrients in Paracou (Schmitt et al. 2021). Waterlogging and topography have been highlighted as crucial for forest dynamics (Ferry et al. 2010), species-habitat relationships (Engelbrecht et al. 2007), and phenotypic variation (Schmitt et al. 2020). TWI was derived from a 1-m-resolution digital elevation model using SAGA-GIS (Conrad et al. 2015) based on a LiDAR campaign of the whole Paracou field station done in 2015.

We also tested the link between functional traits and species growth using the mean trait values of the 120 species that Vleminckx et al. (2021) shared with our study (over 138, Fig. S10). Vleminckx et al. (2021) functional data included 19 leaf, stem and root traits. Leaf traits included carbon, nitrogen, potassium, phosphorus, calcium, and chlorophyll content, carbon 13 isotopic ratio, toughness, thickness, area and specific area. Stem traits included sapwood specific gravity and trunk bark thickness. Root traits included specific tip abundance, specific length, wood specific gravity and fine roots tissue density and diameter.

### Analyses

To study the effect of phylogeny and environment, we investigated the effects of family, genus, species, topography (TWI) and neighbourhood (NCI) indices on individual growth potential (Gmax) with a linear mixed model (model 2). Environmentals variables (TWI and NCI) were used as fixed effects, while taxonomic variables (family, genus, and species) were used as random effects. We propagated the uncertainty in the estimation of individual growth (model 1) to the model testing the effect of phylogeny and environment (model 2). To do so, we considered the individual growth potentials obtained in 500 iterations of model 1 as 500 different dataset on which we fitted a Bayesian linear mixed model (model 2). For each of these dataset, we kept the 1000th iteration after a warm up of 999 iterations. We pooled all these last iterations to obtain a posterior distribution of the effect of phylogeny and environment that takes into account the uncertainty in individual growth potential. We reported the resulting marginal (fixed effects alone) and conditional (fixed and random effects) goodness of fit (R², Nakagawa & Schielzeth 2013).

We further investigated species growth potential across the phylogeny. To account for the lognormal distribution of individual growth potential within species (Schmitt et al. 2022), we used the median as a measure of species growth potential (*Gmax_s_* = *Median*(*Gmax_i_*)). We tested the phylogenetic signal of species growth potential with multiple measures including Pagel’s λ (Keck et *al.*, 2016). We further computed the phylogenetic correlogram of species growth potential using partistic phylogenetic distance and tested its significance with 999 bootstrap assuming brownian motion. We explored the phylogenetic signal structure across the phylogeny with the local indicator of phylogenetic association.

We finally explored the link between species functional traits and species growth potential. We used a linear model with a step selection with stepwise direction of the best model explaining species log-transformed growth potential with log-transformed functional traits to meet the normality assumptions (model 3). We measured functional traits relative importance in explaining species growth potential after stepwise selection using the lmg metric. We used the R^2^ contribution averaged over orderings of regressors as in (Lindeman, Merenda and Gold 1980), using the function *relimp* from the *relaimpo* R package with default parameters (Groemping 2007). We finally compared the taxonomic and phylogenetic structures of functional traits with those of species growth potential using linear mixed models with random taxonomic levels (family, genus, and species) and phylogenetic correlograms. We used the *tidyverse* (Wickham *et al.,* 2019), *lme4* (Bates *et al.,* 2015), *phylosignal* (Keck *et al.,* 2016), and *ggtree* (Yu 2020) packages in the R environment (R Core Team, 2020) for all analyses.

## Results

We recorded 117,688 diameter measurements in the local community across 7,961 individuals belonging to 138 species, 95 genera and 38 families. Bayesian inference of the individual growth model converged correctly (*e.g.,* Fig. S1) with a good mixture of chains for most of the individual growth potentials (*Gmax_i_*) shown with an 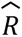 < 1. 1 (99.2%, Fig. S2). The posteriors of the parameters did not show any correlation problems (*e.g.,* Fig. S3). Posterior distributions of individual growth potentials showed limited uncertainty (*e.g.,* Fig. S4). The resulting predicted diameters showed very good goodness of fit with a root mean square error of 0.43 centimetres (Fig. 1, illustrated for 90 individuals in 9 species in Fig. S5).

Based on the inferred individual growth potentials, we found most of the variation in growth potential to be among individuals within species shown with residuals (σ = 0. 72, Tab. 1), then among genera within families (σ = 0. 12), then among species within genera (σ = 0. 04) and among families (σ = 0. 03). The taxonomic structure explained about a third of the observed variation in individual growth potential (Conditional *R*^2^ =0. 347 against Marginal *R*^2^ = 0. 019). The neighbourhood crowding index (NCI) had a marked negative significant effect on individual growth potential (β =− 0. 51, *CI* = [− 0. 61; − 0. 42], Tab. 1, Fig. 2A, Fig. S6) which explained 2% of the observed variation knowing taxonomy (Marginal *R* = 0. 019, Tab. 1). The topographic wetness index had no effect on individual growth potential (β =− 0. 03, *CI* = [− 0. 13; – 0. 07], Tab. 1, Fig. S6).

**Figure 2:**
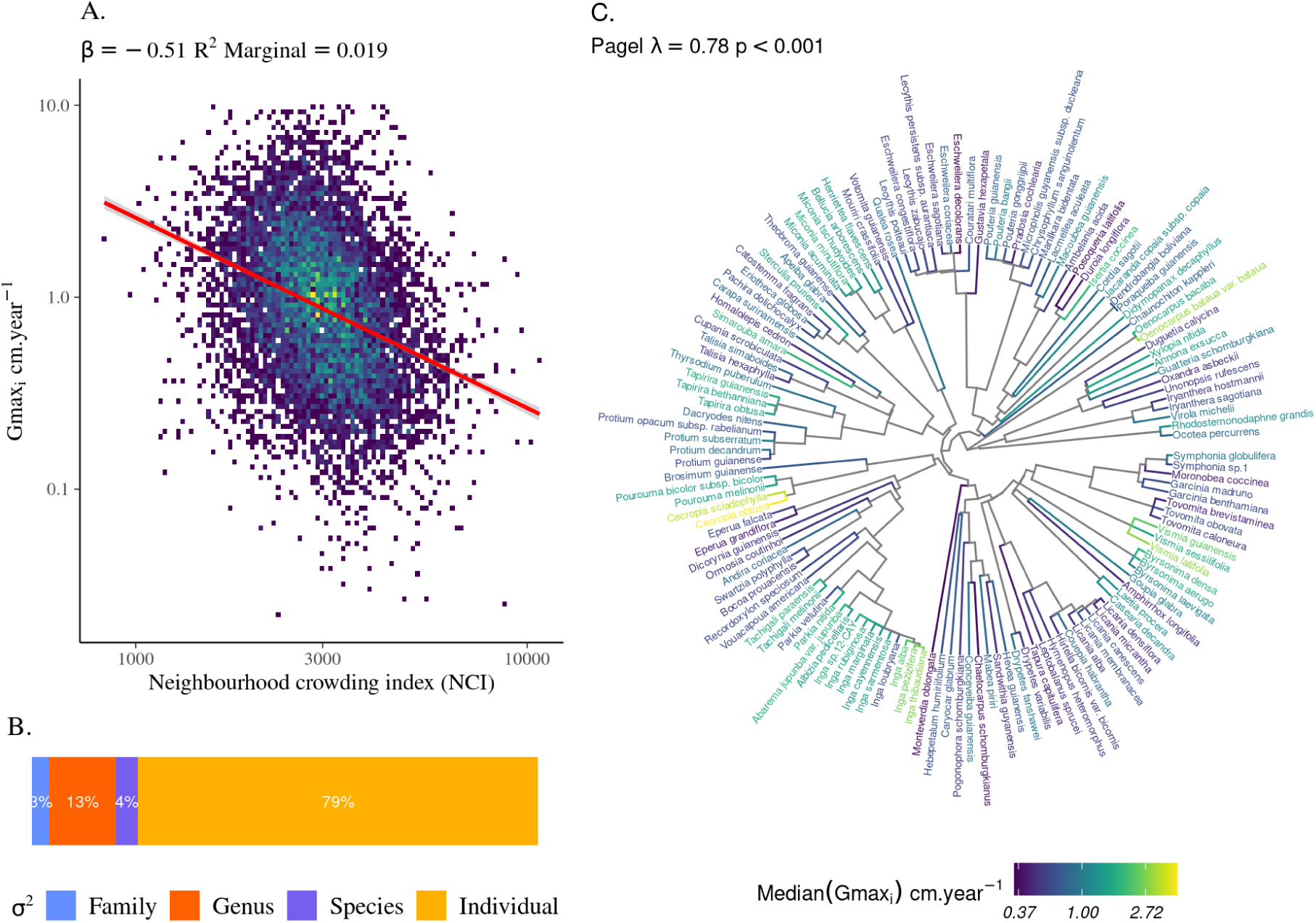
Variation in the growth potential of individuals and species as a function of neighbourhood crowding, taxonomic levels or across phylogeny. **A.** Individual growth potential (Gmax_i_) is significantly decreasing with neighbourhood crowding index (NCI, β =− 0. 51, Marginal R^2^ = 0. 019, see Tab. 1). **B.** The partitioning of the variation of individual growth potential (Gmax_i_) across taxonomy shows that most of the variation occurs at the individual (σ = 0. 72, Tab. 1), then genus (σ = 0. 12) before species (σ = 0. 04) and family (σ = 0. 03) levels. **C.** The distribution of species growth potential (Median[Gmax_i_]) across the phylogeny from slow growing species in dark blue to fast growing species in yellow (log-scale) is phylogenetically structured (Pagel’s λ = 0. 78, p < 0. 001) with a significant positive autocorrelation to a phylogenetic partistic distance below 100 (Fig. S1), corresponding to the genus level (Tab. 1).

**Table 1:** Effects of phylogeny and environment on individual growth potential. We investigated the effects of family, genus, species, topography (TWI) and neighbourhood crowding (NCI) indices on individual growth potential (Gmax) with a linear mixed model. First rows show estimates and credible intervals (CI) for environmental fixed effects. Middle rows show estimates for variance components from phylogenetic random effects. The last row shows marginal (fixed effects alone) and conditional (fixed and random effects) goodness of fit.

| $\log(Gmax_i)$ | | |
| --- | --- | --- |
| *Predictors* | *Estimates* | *CI (95%)* |
| Intercept | 3.88 | 3.05 – 4.68 |
| $\log(NCI_i)$ | -0.51 | -0.61 – -0.42 |
| $\log(TWI_i + 1)$ | -0.03 | -0.13 – 0.07 |
| **Random effects** |  |  |
| Residual (Individual) | 0.72 |  |
| Species \| Genus \| Family | 0.04 |  |
| Genus \| Family | 0.12 |  |
| Family | 0.03 |  |
| Marginal R<sup>2</sup> / Conditional R<sup>2</sup> | 0.019 / 0.347 |  |

Species growth potential was significantly structured in the phylogeny (Pagel’s λ = 0. 78, *p* < 0. 001, Fig. 2B, Tab. S3). Phylogenetic autocorrelograms revealed a short distance significant positive association and a long distant significant negative association of species growth potential in the phylogeny (Fig. S7). The local indicator of phylogenetic association highlighted the conservation of species growth potential at the genus level (Fig. S8), as illustrated for instance with fast growing species from the *Cecropia* genus opposed to slow growing species from the *Eschweilera* genus (Fig. 2C). However, a few species have different growth potential in the same genus, such as slow growing *Drypetes variabilis* opposed to fast growing *Drypetes fanshawei*.

Functional traits explained 40% of the observed variation of species growth potential among 120 tropical tree species (*R*^2^ = 0.394). Leaf potassium content, sapwood specific gravity, fine roots diameter and specific length had significant negative effects on species growth potential, while leaf nitrogen content had a significant positive effect (Tab. 2). Sapwood specific gravity and leaf nitrogen content had the largest relative importance (resp. *R*^2^ = 0. 18 and *R*^2^ = 0. 11), while other traits had small relative importance (*R*^2^ ≤ 0. 04). Moreover, these six functional traits had a phylogenetic autocorrelogram similar to species growth potential with a short distance significant positive association, stronger for sapwood specific gravity than other traits (Fig. S9).

**Table 2:** Effects of functional traits on species growth potential. We extracted traits from Vleminckx et al. (2021) for the 120 species shared with our study, and used a linear model with a step selection of the best model explaining species growth potential with log-transformed growth potential and functional traits. We thus selected 6 root, stem and leaf functional traits. Columns show estimates, confidence intervals (CI), significance (p-value), and relative importance (R²) for selected functional traits. The last row shows raw and adjusted goodness of fit.

| $\log(Gmax_s)$ | | | | |
| --- | --- | --- | --- | --- |
| *Predictors* | *Estimates* | *CI* | *p-value* | *R<sup>2</sup>* |
| Intercept | -0.01 | -0.15 – 0.13 | 0.882 |  |
| leaf toughness | 0.18 | -0.01 – 0.37 | 0.063 | 0.02 |
| leaf nitrogen content | 0.38 | 0.22 – 0.55 | **<0.001** | 0.11 |
| leaf potassium content | -0.24 | -0.41 – -0.07 | **0.005** | 0.04 |
| sapwood specific gravity | -0.57 | -0.76 – -0.39 | **<0.001** | 0.18 |
| fine roots diameter | -0.49 | -0.82 – -0.17 | **0.003** | 0.04 |
| specific root length | -0.35 | -0.67 – -0.02 | **0.035** | 0.03 |
| R<sup>2</sup> / R<sup>2</sup> adjusted | 0.424 / 0.394 |  |  |  |

## Discussion

Using 36 years of diameter records for thousands of mapped individuals belonging to 138 species, we found that the evolutionary heritage of species shaped individual growth trajectories, explaining up to one third of the observed variation in growth potential among the eight thousand individuals studied. Functional traits of wood, leaves and roots together predicted species growth potential, showing that multiple functional dimensions determine the performance of tropical tree species in their environments. Phylogenetic correlograms suggested joint selection of species’ growth strategies and associated functional traits during convergent evolutions. Nevertheless, at the individual tree level, the observed effect of neighbourhood crowding suggests that forest gap resulted in fast-growing trees in high light conditions with reduced competition, as opposed to slow-growing trees in low light conditions with strong competition. But the ecological and evolutionary drivers of the high variability in individual growth potential remain largely undetermined, and the underlying factors, which include phenotypic plasticity and genetic adaptations, are little explored. The high intraspecific variation observed could allow individuals in hyperdiverse ecosystems such as tropical forests to respond to the variable light and competitive conditions offered by successional niches during forest gap dynamics, in addition to the many other potential environmental dimensions that shape the coexistence of species.

### Evolutionary history shapes the growth of tropical trees

We found that evolutionary history shaped individual growth potential. Previous studies already highlighted regionally a phylogenetic signal in species growth potential in the Amazon Basin, with evolutionarily related genera having more similar growth values than expected by chance (de Souza *et al*., 2016). Regionally, woody biomass and tree size also showed a strong phylogenetic signal in tropical dry forests (de Aguiar-Campos *et al*., 2021), besides longevity rather than growth determined biomass (Körner 2017). Similarly, Cadotte *et al.,* (2009) found evolutionary relationships explaining grasslands productivity. Thus, evolutionary history shapes observed phylogenetic and functional diversity at the regional scale, with for instance lasting influences from the Paleoclimate (Svenning *et al*., 2015; Bosela *et al*., 2016). Our results are in agreement as we also identified a strong phylogenetic signal in species growth potential with positive phylogenetic correlation up to the genus level, showing that previous regional scale results hold locally. To our knowledge, our study is the first attempt to link individual growth potential to species growth potential. Using taxonomy as a proxy for evolutionary heritage of species, we found that the evolutionary history of species explained up to one third of the observed variation in growth potential among the eight thousand individuals studied. Thus, evolutionary history is an important determinant of individual variation in growth that is stronger than the direct effect of the environment, at least for the topography and neighbourhood crowding tested here.

Several convergent evolution events resulted in similar growth patterns of species within genera with, for instance, repeated evolution of fast-growing species in *Urticaceae*, *Fabaceae*, *Hypericaceae*, and *Melastomataceae* families with species respectively belonging to the *Cecropia*, *Inga*, *Vismia* and *Miconia* genera. This pattern might be explained by repeated evolution constrained by forest gap dynamics, leading to shade-tolerance under closed canopies as opposed to fast-growing pioneer species in light gaps (de Souza *et al*., 2016). Indeed, *Cecropia, Visimia* and *Miconia* are for instance recognized as pioneer species colonising first light gaps after a treefall (Dalling *et al*., 1998). Convergent evolutions of habitat specialisation could also explain the divergent growth of genera within families. For instance at the species level, Fine *et al.,* (2014) evidenced specialisation to white sands or flooded soils within *Protieae*. Nevertheless, the fact that topography and forest gap dynamics, two major determinants of the local community (e.g. Ferry *et al.,* 2010; Molino and Sabatier 2001), have less explanatory power than taxonomy reveals the importance of past evolutionary constraints on the growth trajectories of individual trees through species heritage compared to the direct effect of the current environment to which individual trees could respond through microadaptations and phenotypic plasticity (*e.g.* Schmitt *et al*., 2021; 2022).

Taken together, our results therefore partly support species-averaged community ecology approaches, at least for growth trajectories (but see last paragraph). Swenson *et al*., (2013) suggested the use of phylogenetic approaches to understand community assembly. However, phylogeny alone is not sufficient to predict demographic rates (Che-Castaldo *et al*., 2018). Indeed, phylogenetic approaches must be conducted with caution given the high remaining intraspecific variability (see last paragraph) and the differences observed among closely-related species.

### Multiple functional dimensions together predict the species growth potential

Using the 120 species in common with Vleminckx *et al*. (2021), we found six functional traits explaining an important part of variation in species growth potential (40%). Sapwood specific gravity was already shown to be a major predictor of species growth (King et al., 2005; King et al., 2006; Hérault et al., 2011; Visser et al., 2016), with fast-growing species investing less in wood resulting in a low density as opposed to slow-growing ones. Leaf nitrogen content was the second most important predictor in our study, with nitrogen-rich species growing faster, as already observed for pioneer species (Aidar *et al*., 2003), and in rich environments (Russo et al., 2008). Leaf potassium and toughness were also small predictors of species growth potential. Finally, root traits also predicted species growth potential with fast-growing species investing less in their root systems with decreased fine roots diameters and higher specific root length, as already shown for temperate species (Comas, Bouma and Eissenstat 2002, Comas and Eissenstat 2004) but never in the tropics, in the limit of our knowledge. Soil heterogeneity can be expected to play a key role in shaping root traits (Vleminckx *et al.,* 2021), while the dynamics of forest gaps cannot be excluded either (Xiang *et al*., 2013).

Future research on the role of root traits in tropical tree growth is thus particularly promising given that tropical forests harbour the greatest diversity of root characteristics (Ma *et al*., 2018). Interestingly, species growth potential is better predicted by the combination of traits from roots, wood and leaf, in accordance with the idea of multiple functional dimensions that allow tropical tree species to optimise their performance in a given environment Vleminckx *et al*. (2021), while participating together in a whole plant economic spectrum (Reich 2014).

Finally, we found a similar phylogenetic structure between the 6 functional traits and species growth potential, with significant and positive phylogenetic correlation at short distance, in particular in sapwood specific gravity (de Souza *et al*., 2016). The traits explaining the most species growth had the strongest positive phylogenetic correlation (*e.g.* sapwood specific gravity), while traits such as leaf potassium content and toughness had smaller positive phylogenetic correlation only significant to a short phylogenetic distance. The similarity between phylogenetic correlograms suggests a joint selection of species growth strategies and associated functional traits during convergent evolutions. This result further supports the use of phylogeny and related traits to predict species growth trajectories and potential demography (Swenson 2014; Tucker *et al*., 2018; Paquette *et al.,* 2015).

### Individual growth potential is influenced by forest gap dynamics but remains largely unexplained

We found evolutionary heritage shaping the individual growth of tropical trees, with functional traits as important predictors of species growth potential. However, we still observe a huge intraspecific variation with individual growth potential varying widely within species (logarithmic coefficient of variation of 98% [35-430%]). The growth trajectories of individual trees are strongly and negatively affected by the average neighbourhood crowding over the last 30 years, which can be related to the mosaic of light and competition environments shaped by forest gap dynamics (Schmitt *et al.,* 2022). In a nutshell, we observe fast-growing trees in high-light conditions with decreased competition opposed to slow-growing trees in low-light conditions with strong competition within and among species. Fast-growing pioneer species opposed to slow-growing species under closed-canopies are widely known and expected at the interspecific level (Dalling*et al*., 1998; Molino and Sabatier 2001; King et al., 2005; Hérault et al., 2011), but we also evidenced fast-growing individual opposed to slow-growing individual within species along successional niches, advocating for a wide breadth of successional niches in tropical tree species (e.g. Schmitt *et al.,* 2022 for *Symphonia* species). The effect of competition and its reduction has already been suggested as a factor increasing tree radial growth within species in the context of selective logging, especially within shade-tolerant species (Peña-Claros *et al.,* 2008). Indeed, logging gaps, with increased light access, results in increased tree growth at short distance from the gap, especially within slow-growing species (Hérault *et al.,* 2010). Rare species have been also shown to be more sensitive to light variation (Rüger *et al.,* 2011). The effect of crowding can be a direct limit to light access expected in successional niches, but the effect of neighbours identity found in tropical trees (Potvin *et al.,* 2008) also suggest other above- and below-ground competition and facilitation processes. In short, forest gap dynamics with tree fall results in successional niches ranging from high light and low competition after recent tree fall to low light and high competition in closed canopies. We found along this gradient fast-growing species and individuals within species in early-succession niches as opposed to slow-growing species and individuals within species in late-succession niches.

Nevertheless, forest gap dynamics only explained a tenth of the high variation of individual growth potential within species. Topographic position, proxied with topographic wetness index, did not influence the individual growth variation within species in our study (but see O’Brien and Escudero 2021), despite its known importance in tropical forest dynamics (Ferry *et al.,* 2010). The role of topography on individual growth potential was weak, but we can assume that topography shapes species growth through species evolutionary heritage, as several species studied showed locally pervasive habitat preferences along topography (Allié *et al.,* 2018; Schmitt *et al.,* 2021b) and microgeographic adaptations to topography (Schmitt *et al.,* 2021). Consequently, ecological and evolutionary factors of individual growth potential remain largely undetermined. The process through which forest gap dynamics and undetermined factors affect individual growth potential also remains underexplored. The spatio-temporal variation of forest gap dynamics led to assume growth potential variation within species to be due to phenotypic plasticity (dos Santos *et al.,* 2020). However, recent studies revealed local adaptation (O’Brien and Escudero 2021) and microadaptation (Schmitt *et al.,* 2022) of individual trees within species to neighbourhood crowding and competition resulting in varying individual growth potential. Thus forest gap dynamics could have both a strong evolutionary heritage on species and may shape genotypes within species with strong spatio-temporal variations (Schmitt *et al.,* 2022).

Intraspecific variability in performance can have strong implications for species coexistence (Chesson 2000, Clark 2010, see also Stump *et al.,* 2022 for a synthesis), although this view remains an open question (*e.g.* Clark *et al.,* 2022). Clark (2010) suggested that intraspecific variability allows species to differ in the distribution of their responses to the environment and thus to pass environmental filtering: an individual may persist in a given environment with a suitable phenotype while the same environment would have filtered out the average phenotype of the species. This hypothesis is consistent with theories that predict the coexistence of a greater number of species, with competition being stronger within species, among individuals, than among species (Chesson 2000). Modelling approaches further support the hypothesis that intraspecific genetic and phenotypic variability promotes species coexistence (Lichstein *et al.,* 2007). In the case of forest gap dynamics, late-successional species have been shown to have more variation in response to competition and light variation than early-successional species (Peña-Claros *et al.,* 2008; Hérault *et al.,* 2010), which could be linked to a greater diversity of light and competitive environments. The high intraspecific variation observed (logarithmic coefficient of variation of 98%) could therefore allow individuals in these hyperdiverse ecosystems to adjust to the variable conditions of light and competition offered by the successional niches during the closure of the forest gaps, in addition to the other numerous potential niches shaping the high-dimensional coexistence of species (Clark 2010). Consequently, the methodology used in our study paves the way to future research on determinants and processes shaping tree growth within and among species. The combination of studies from forest censuses (e.g. this study), modelling approaches (e.g. Schmitt et al., 2020b), and experimental studies (e.g. O’Brien and Escudero 2021) holds promise for a better understanding of tree performance within and among species, with the potential to better explain and predict species coexistence and forest dynamics. In particular, linking individual genotypes within species to forest gap dynamics, topography and individual growth potential (Schmitt *et al.,* 2022) among several species to extrapolate to the community level seems a promising approach to elucidate the determinants of tree growth.

## Acknowledgements

We are grateful to Pascal Petronelli, Julien Engel, Christopher Baraloto and the CIRAD inventory team for their tremendous work in tree inventory and botanical identification, without which this entire study would simply not have been possible.

## Data, script and code availability

DBH and spatial positions of individuals were extracted from the Paracou Station database, for which access is available at https://dataverse.cirad.fr/dataverse/paracou with corresponding DOIs given in table S4. Scripts used for analyses can be found on GitHub (https://github.com/sylvainschmitt/treegrowth).

## Authors contribution

SS conceived the ideas and designed the methodology. SS, BH, and GD analysed model outputs. SS led the writing of the manuscript. All authors contributed critically to the drafts and gave final approval for publication.

