## Supplementary material for "High intraspecific growth variability despite strong evolutionary heritage in a neotropical forest": Fig. S1

The following Supporting Information is available for this article:

**Tab. S1.** Model shape selection.

**Tab. S2.** Model selection for hierarchical integration of individual and species effects.

**Tab. S3.** Phylogenetic signals for species growth potential.

**Tab. S4.** Digital Object Identifiers (DOI) for used Paracou datasets.

**Fig. S1.** Traceplot of chains in the individual growth model.

**Fig. S2.** Convergence of parameters in the individual growth model.

**Fig. S3.** Pairs plots of parameters in the individual growth model.

**Fig. S4.** Posterior distribution 100 individual growth potentials.

**Fig. S5.** Predicted diameter trajectories for 90 individuals in 9 species.

**Fig. S6.** Effects of the environment on individual growth potential.

**Fig. S7.** Phylogenetic autocorrelogram of species growth potential.

**Fig. S8.** Local indicator of phylogenetic association of species growth potential.

**Fig. S9.** Phylogenetic autocorrelogram of species growth potential and functional traits.

*Table S1: Model shape selection based on quality of prediction (Root Mean Square Error of Prediction, RMSEP, cm), cross-validation (Leave-One-Out Estimate of the expected Log pointwise Predictive Density, LOO ELDP), and speed (Elapsed time, seconds). The selected model is shown in bold.*

| **Model** | **Equation** | **RM SEP** | **LOO EDLP** | **Elapsed time** |
|---|---|---|---|---|
| Michaelis Menten | $DBH_t \sim N(DBH0 + \frac{\alpha \times t}{\beta + t}, \sigma)$ | 6.8 | 214 | 5 |
| Gompertz | $DBH_t \sim N(DBH0 + AGR_{t-1}, \sigma)$ | 0.4 | 253 | 51 |
| Lognormal | $DBH_t \sim N(DBH0 + \beta \times log(t), \sigma)$ | 1.6 | 224 | 2 |
| Polynomial | $DBH_t \sim N(DBH0 + \alpha \times t + \beta \times t^2 + \gamma \times t^3, \sigma)$ | 0.3 | 228 | 40 |
| Weibull | $DBH_t \sim N(DBH0 + \alpha \times (1 - e^{-t \backslash \beta}), \sigma)$ | 0.8 | 218 | 6 |
| Amani | $DBH_t \sim N(DBH0 + \alpha \times (1 - e^{-\lambda(\frac{t}{\theta})^{\beta}}), \sigma)$ | 6.3 | 243 | 100 |
| **Gompertz sum** | $DBH_t \sim N(DBH0 + \sum_{y=0}^{y=t} AGR_y, \sigma)$ | **0.4** | **248** | **208** |

*Table S2: Model selection for hierarchical integration of individual and species effects on optimal diameter (Dopt) and kurtosis (Ks) based on quality of prediction (Root Mean Square Error of Prediction, RMSEP, cm), cross-validation (Leave-One-Out Estimate of the expected Log pointwise Predictive Density, LOO ELDP), and speed (Elapsed time, seconds). The selected model is shown in bold.*

| **Model** | **RMSEP** | **LOO EDLP** | **Elapsed time** |
|---|---|---|---|
| Individual fixed effect | 6.8 | 214 | 5 |
| **Individual random effect centred on species fixed effect** | **0.4** | **253** | **51** |
| Species fixed effect | 1.6 | 224 | 2 |
| Species random effect centred on community mean value | 0.3 | 228 | 40 |

*Table S3: Phylogenetic signals for species growth potential with several indices. The p-value is given in brackets.*

| **Indice** | **species growth potential** |
|---|---|
| Abouheif's C | **0.4373 (p=0.001)** |
| Gittleman's I | **0.1456 (p=0.001)** |
| Blomberg's K | 0.0799 (p=0.074) |
| Blomberg's K* | 0.0845 (p=0.077) |
| Pagel's λ | **0.7757 (p=0.001)** |

*Table S4: Digital Object Identifiers (DOI) for used Paracou datasets. Data can be accessed and retrieved at https://dataverse.cirad.fr/dataverse/paracou. Please use "contact the owner" to ask for data access. Reference datasets as Derroire, Géraldine; Hérault, Bruno; Rossi, Vivien; Blanc, Lilian; Gourlet-Fleury, Sylvie; Schmitt, Laurent, 2022, {Name}, {DOI}, CIRAD Dataverse, V1.*

| **Name** | **DOI** |
|---|---|
| Paracou Biodiversity Plots | https://doi.org/10.18167/DVN1/NSCWF0 |
| Paracou Disturbance Experiment - Control Plots | https://doi.org/10.18167/DVN1/Q8V2YI |
| Paracou Disturbance Experiment - Level1 Treatment Plots | https://doi.org/10.18167/DVN1/LIVCEK |
| Paracou Disturbance Experiment - Level2 Treatment Plots | https://doi.org/10.18167/DVN1/HWTD4U |
| Paracou Disturbance Experiment - Level3 Treatment Plots | https://doi.org/10.18167/DVN1/HIGNWQ |

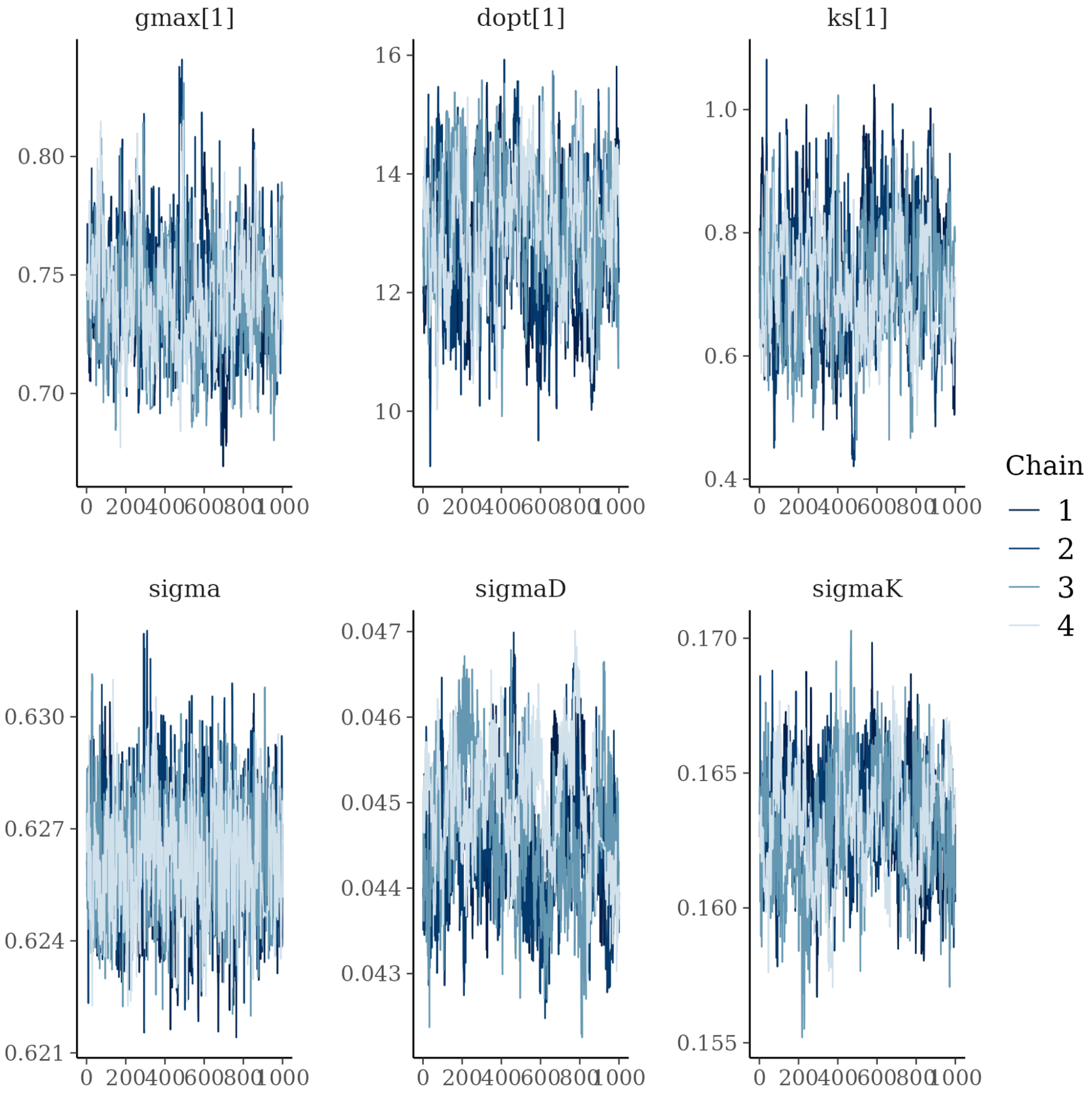


*Figure S1: Traceplot of chains in the Bayesian inference of the individual growth model for 6 parameters: individual growth potential (gmax), optimal diameter (dopt) and kurtosis (ks) for the first individual and associated variances sigma, sigmaD and sigmaK.*

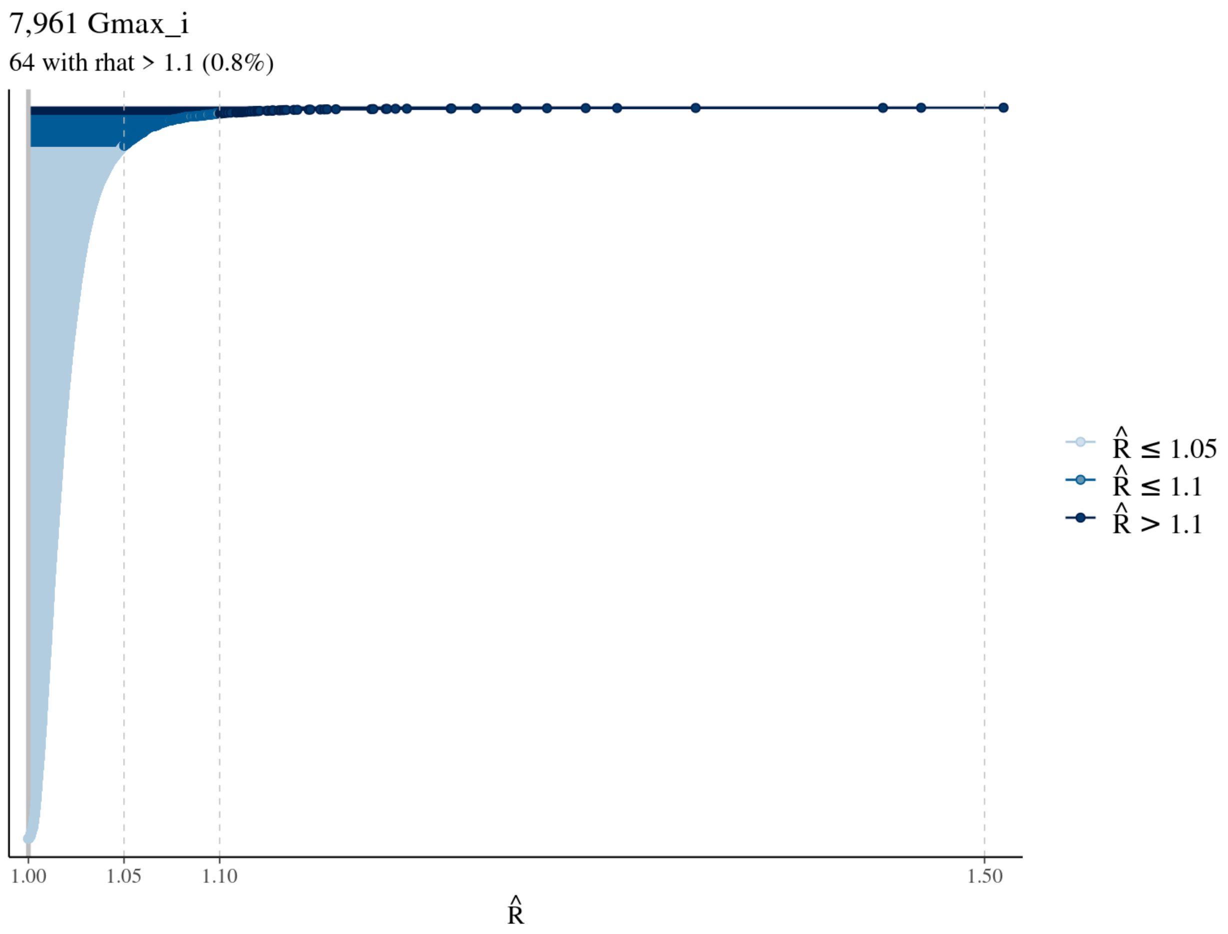


*Figure S2: Convergence of individual growth potential in the Bayesian inference of the individual growth model for the 7,961 individual. A* $\widehat{R} < 1.1$ *indicates a good mixing of chains, and thus a correct convergence of the model.*

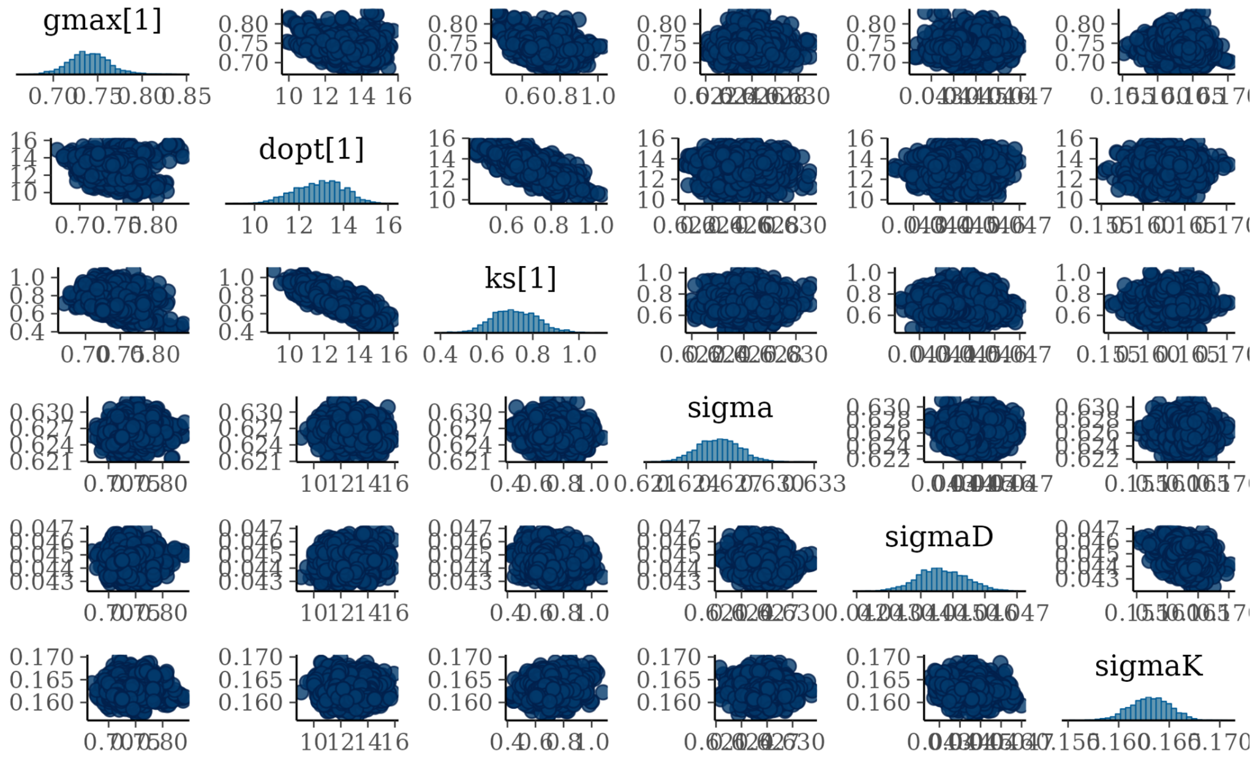


*Figure S3: Pairs plots of parameters values in the Bayesian inference of the individual growth model for 6 parameters: individual growth potential (gmax), optimal diameter (dopt) and kurtosis (ks) for the first individual and associated variances sigma, sigmaD and sigmaK.*

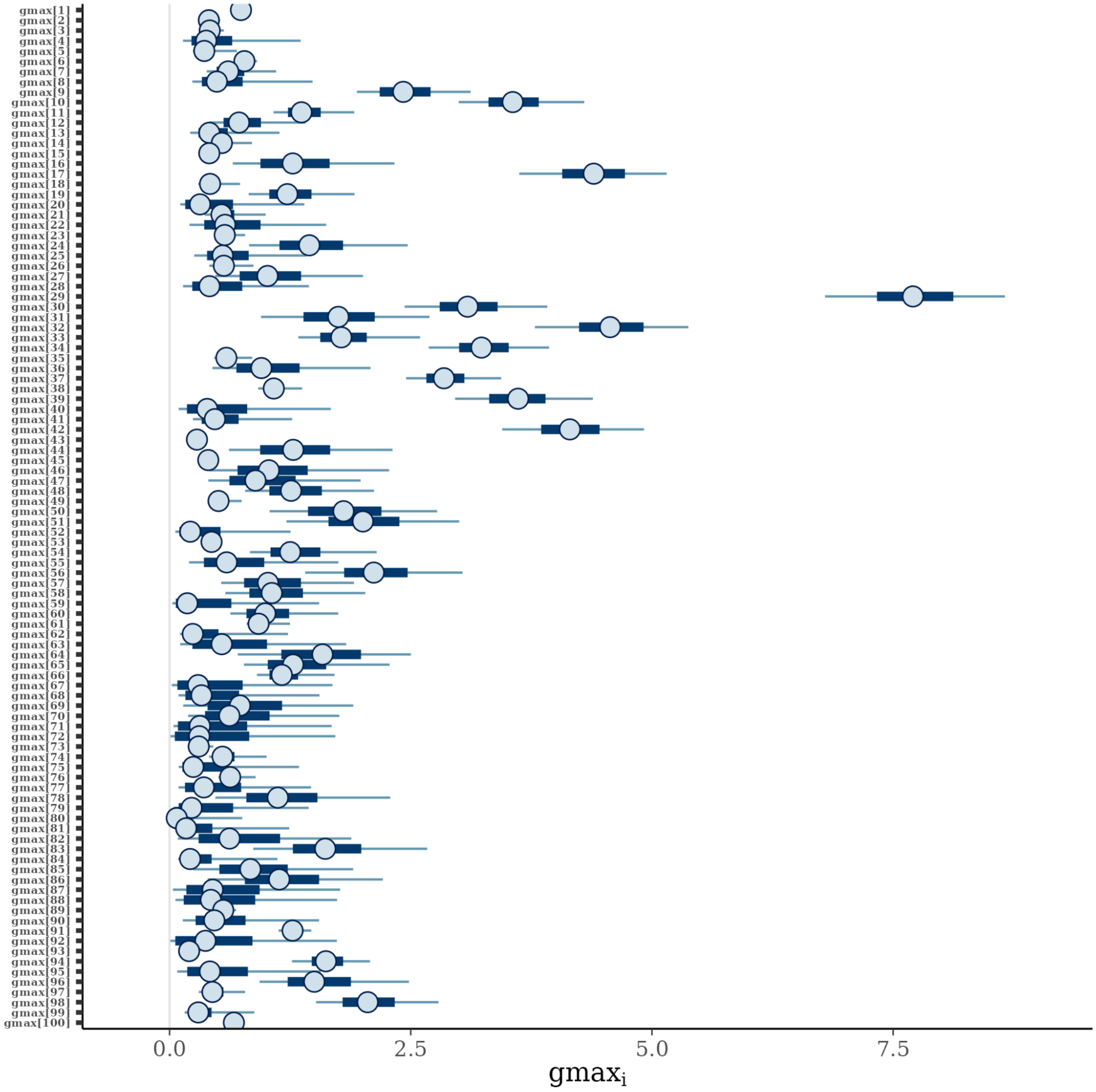


*Figure S4: Posterior distribution of the individual growth potential for the first hundred individuals in the Bayesian inference of the individual growth model. The circle represents the median value, whereas the thick line represents the 50% credibility interval and the thin line the 95% credibility interval.*

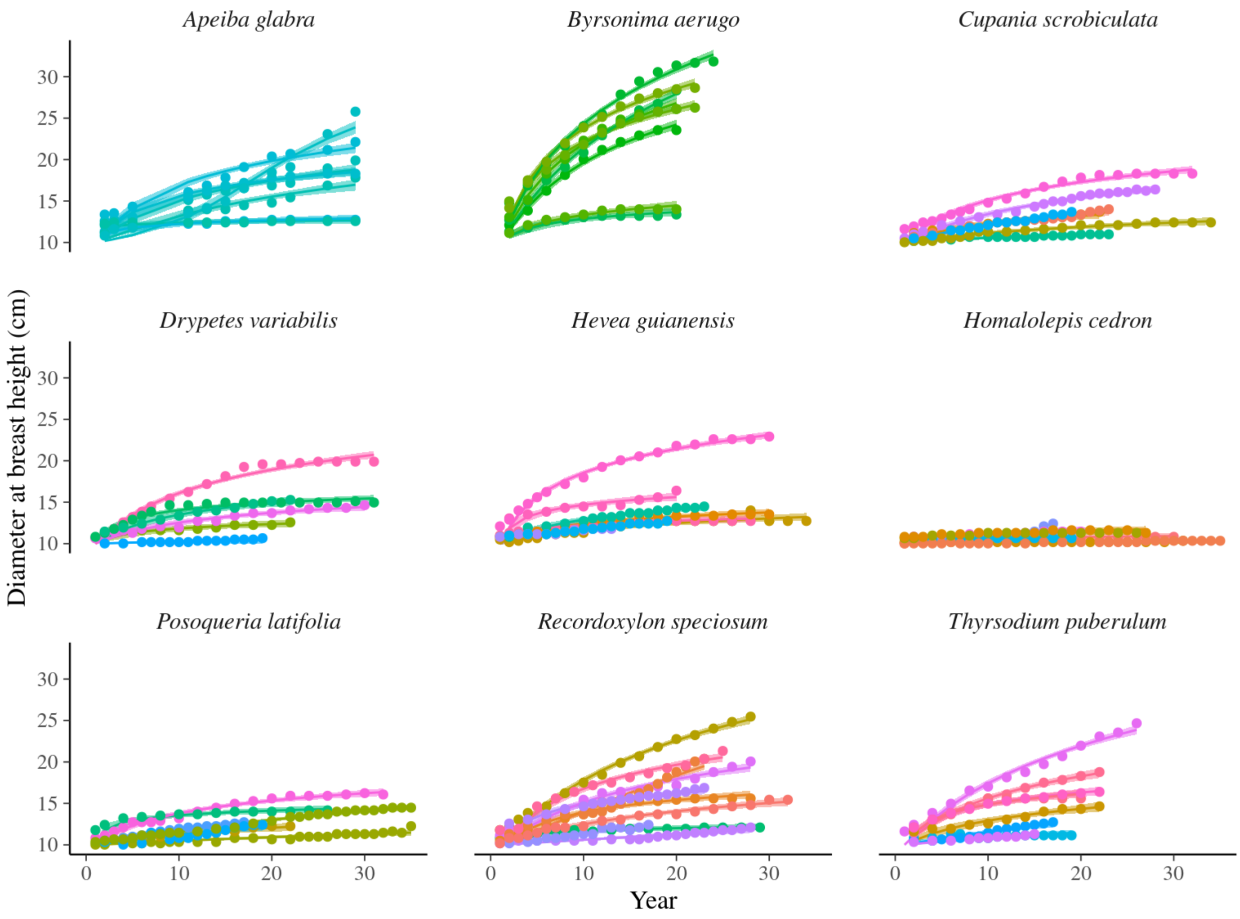


*Figure S5: Predicted diameter trajectories for 90 individuals in 9 species compared to observed diameters at breast height. The points indicate observed diameters whereas the lines represent the medians of the predicted trajectories and the shaded areas the 90% credibility interval. The colours help decipher the individuals in each species.*

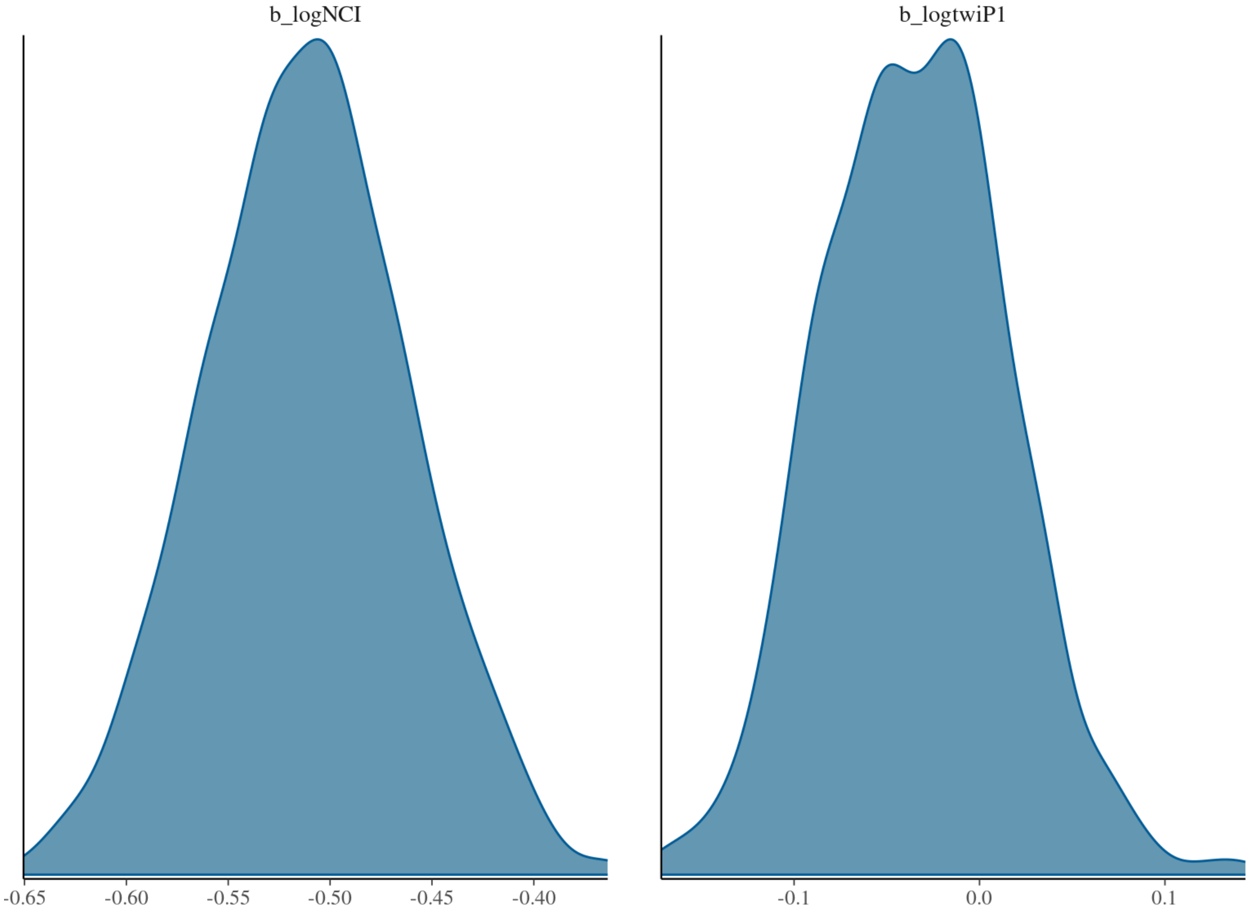


*Figure S6: Posterior distributions for the effects of the environment on individual growth potential with uncertainty transmission. b_logNCI represents the posterior of the effect of neighbourhood crowding index (NCI) on individual growth potential and b_logtwiP1 the effect of topographic wetness index (TWI). The distributions reveal a null effect of TWI but a significantly negative effect of NCI. The distributions account for the uncertainty in individual growth potential (see material and methods).*

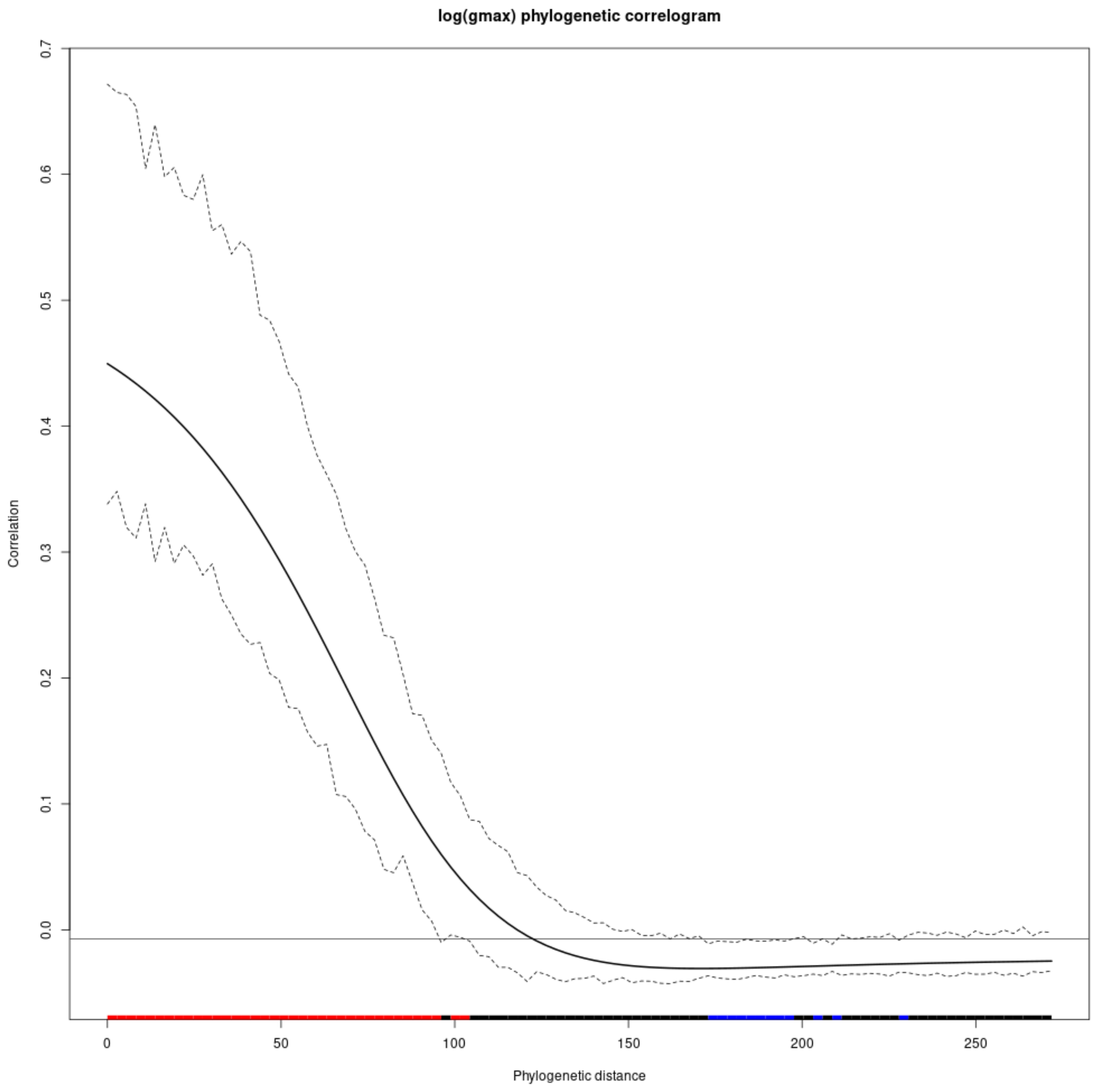


*Figure S7: Phylogenetic autocorrelogram of species growth potential. Significance has been tested with 999 bootstrap assuming brownian motion, with significant positive association in red and a significant negative association in blue.*

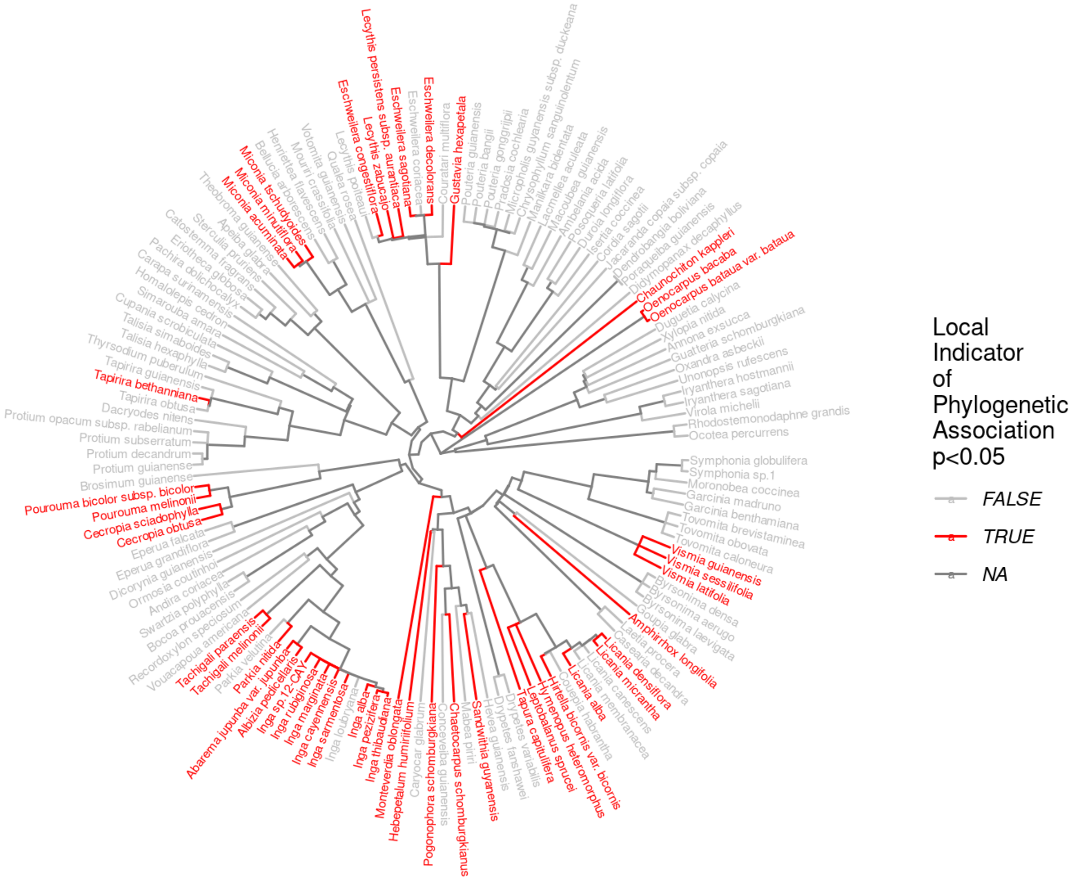


*Figure S8: Local indicator of phylogenetic association of species growth potential. The significance test only concerns the species level (end of the branches), the NA values concern the rest of the branches which are not concerned by the significance test.*

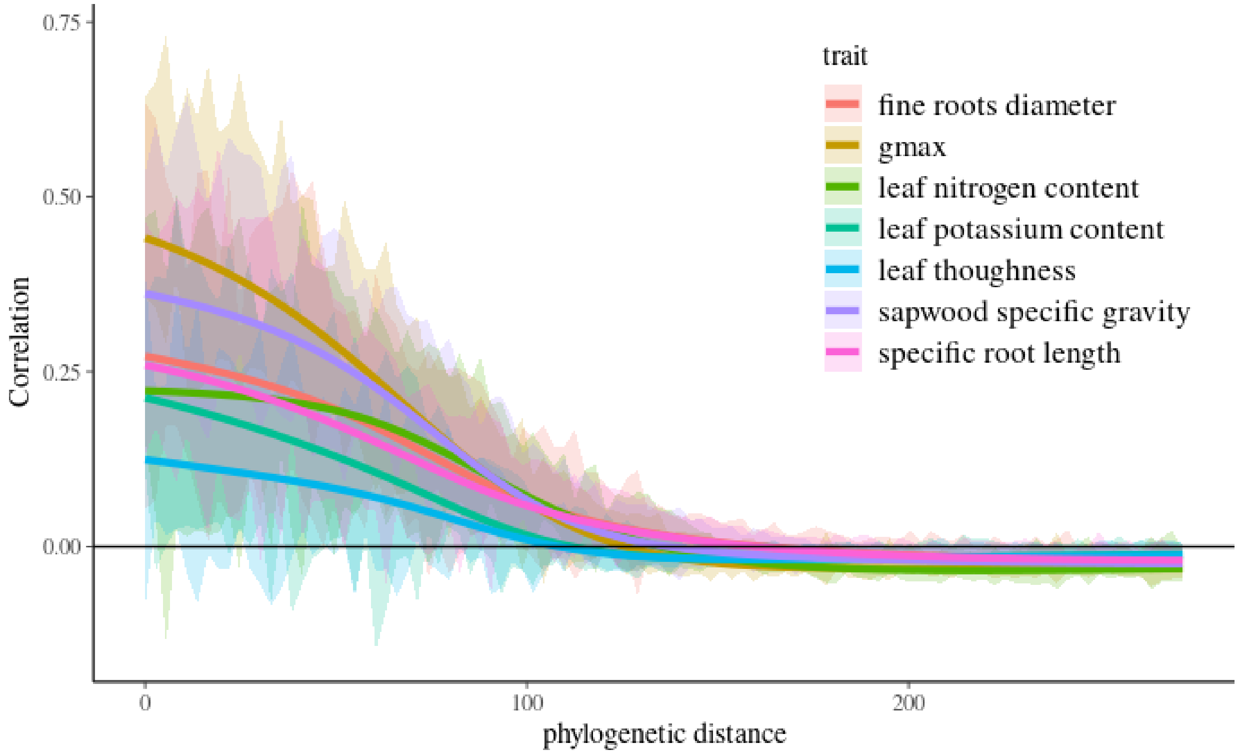


*Figure S9: Phylogenetic autocorrelogram of species growth potential and selected functional traits explaining growth potential. Significance has been tested with 99 bootstrap assuming brownian motion.*

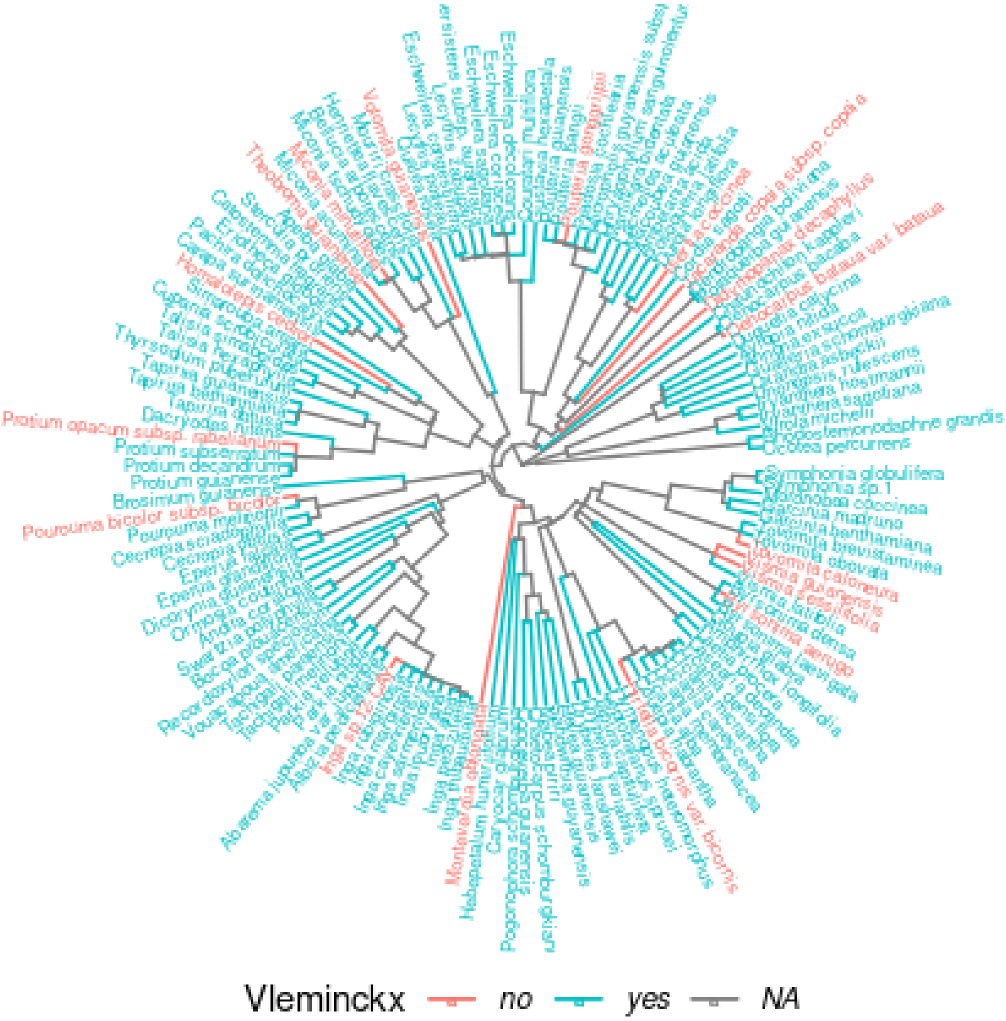


*Figure S10: Distribution of species with functional trait data in the phylogeny. 120 of 138 species are included with no phylogenetic signal of the two groups. The classification only concerns the species level (end of the branches), the NA values concern the rest of the branches which are not concerned.*
